# NAD^+^ prevents septic shock-induced death by non-canonical inflammasome blockade and IL-10 cytokine production in macrophages

**DOI:** 10.1101/2020.03.29.013649

**Authors:** Jasper Iske, Rachid El Fatimy, Yeqi Nian, Amina Ghouzlani, Siawosh K. Eskandari, Hector Rodriguez Cetina Biefer, Anju Vasudevan, Abdallah Elkhal

**Author notes:** These authors contributed equally to this work (Jasper Iske, Rachid El Fatimy, Yeqi Nian). **Corresponding author:** Abdallah Elkhal, Ph.D., Cardiovascular Department, Huntington Medical Research Institutes, Pasadena, CA.

## Abstract

Septic shock is characterized by an excessive inflammatory response depicted in a cytokine storm that results from invasive bacterial, fungi, protozoa, and viral infections. Non-canonical inflammasome activation is crucial in the development of septic shock promoting pyroptosis and pro-inflammatory cytokine production via caspase-11 and Gasdermin-D (GSDMD). Here, we show that NAD^+^ treatment protected mice towards bacterial and LPS induced endotoxic shock by blocking the non-canonical inflammasome specifically. NAD^+^ administration impeded systemic IL-1β and IL-18 production and GSDMD-mediated pyroptosis of macrophages via the IFN-β/STAT-1 signaling machinery. More importantly, NAD^+^ administration not only improved casp-11 KO (knockout) survival but rendered WT mice completely resistant to septic shock via the IL-10 signaling pathway that was independent from the non-canonical inflammasome. Here, we delineated a two-sided effect of NAD^+^ blocking septic shock through a specific inhibition of the non-canonical inflammasome and promoting immune homeostasis via IL-10, underscoring its unique therapeutic potential.

## Introduction

Sepsis is characterized by a systemic inflammatory response syndrome (SIRS)^1^ driven by host cells following systemic bacterial^2^ and viral infections. The excessive inflammatory response can derail into septic shock resulting in multiple organ failure (MOF), the leading cause of death in intensive care units. Inflammasome activation, which downstream pathways cause the release of proinflammatory cytokines and the induction of an inflammatory cell death termed pyroptosis^3^, has been pointed out as the major driver of septic shock. Hereby, a two-armed LPS derived induction of the NLRP3-canonical inflammasome, the major source of IL-1β and IL-18 cytokine production^4^ and the caspase-11 mediated non-canonical inflammasome leading to pyroptosis in monocytes^5^ was determined as the underlying mechanism. Mechanistically, Caspase-11 acts as a pattern recognition receptor for intracellular bacteria^6^ that cleaves gasdermin-D, a membrane-pore forming protein subsequently inducing pyroptotic cell death^7^. The NLRP3-canonical inflammasome in turn, was found to be indispensable^8^ for septic shock induced death. However cross-activation through Caspase-11 promoting cytokine release has been described^9, 10, 11^, assigning the non-canonical inflammasome a cardinal role^12^.

Recent approaches such as anti-proinflammatory cytokine strategies, blocking downstream targets of inflammasomes have been ineffective^13^ while inhibiting inflammatory key regulators such as NF-κB may promote adverse side-effects^14^. Hence, contemporary clinical therapy of septic shock is based on symptomatic treatment rather than curative approaches that clear the cause of the disease itself.

In our previous studies, we have underscored the immunosuppressive properties of NAD^+^ in autoimmune diseases and allo-immunity via the regulation of CD4^+^ T cell fate^15, 16^. More recently, we have shown that NAD^+^ administration protected mice from lethal doses of *Listeria monocytogenes* (*L. m.*) via mast cells (MCs) exclusively and independently of major antigen presenting cells (APCs)^17^. However, the underlying mechanism that allows NAD^+^, to concomitantly protect against autoimmune diseases, via its immunosuppressive properties^15, 16^, and against lethal bacterial infection remains unclear.

Therefore, in the current study we investigated whether NAD^+^ protects against bacterial infection by dampening the systemic inflammatory response associated with sepsis or through enhanced bacterial clearance. Although, wild type (WT) mice subjected to NAD^+^ or PBS and lethal doses of pathogenic *Escherichia coli* (*E. coli*) exhibited similar bacterial load in various tissues, mice treated with NAD^+^ displayed a robust survival. Moreover, NAD^+^ protected against LPS-induced death that was associated with a dramatic decrease of systemic IL-1β and IL-18 levels, two major cytokines involved in the inflammasome signaling machinery. More importantly, we show that NAD^+^ protected from LPS-induced death by targeting specifically the non-canonical inflammasome via a blockade of the STAT-1/IFN-β signaling pathway. Moreover, NAD^+^ treatment rendered not only Caspase-11 KO but WT mice fully resistant to poly(I:C) + LPS induced septic shock, via an inflammasome independent pathway mediated by a systemic IL-10 cytokine production.

## Results

### NAD^+^ protects mice against septic shock not via bacterial clearance but via inflammasome blockade

Our previous studies have underscored the role of NAD^+^ in regulating CD4^+^ T cell fate and its immunosuppressive properties via IL-10 cytokine production^15, 16, 17^. More recently, we have shown that NAD^+^ protected mice against lethal doses of *L. m.* independently of major APCs^15^. However, it remained unclear whether NAD^+^ protected mice against lethal doses of *L. m*., a gram-positive bacterium, via a clearance mechanism or by dampening the inflammatory response. Since *L. m.* is known to be an intracellular pathogen, we tested if NAD^+^ protects as well against *E. coli*, a gram-negative bacterium that is well known to induce septic shock^18^. Wild type mice were treated with NAD^+^ or PBS for 2 consecutive days followed by a lethal dose (1×10^9^) of *E. coli*. or PBS. Notably, mice treated with PBS died within 5 hours after infection, while mice treated with NAD^+^ exhibited an impressive survival (**Fig. 1A**). Moreover, when assessing the bacterial load in liver and kidney (**Fig. 1B**), organs exposed to the infection, by counting CFU in both, NAD^+^ and PBS groups, revealed no significant difference, suggesting that NAD^+^ does not promote bacterial clearance. More importantly, these data suggest that NAD^+^ may reduce the inflammatory response towards bacterial infection. It is well established that the bacterial lipopolysaccharide (LPS) abundant on the outer membrane exhibits a key role in the pathology of *E. coli* derived septic shock^13^. Thus, we further characterize the impact of NAD^+^ on septic shock by subjecting mice to a lethal dose (54mg/kg) of two different LPS serotypes (O111:B4 and O55:B5) described to vary in the antigen lipid A content and to promote distinct hypothermia kinetics^19^. Following LPS (O111:B4 and O55:B5) administration, PBS treated control mice displayed severe symptoms of endotoxic shock with a dramatical body temperature decrease (<23°C) within 15 hours. In contrast, mice subjected to NAD^+^ exhibited highly distinct kinetics with a recovery of body temperatures after 15 hours (**Fig. 1C**). When monitoring survival, 100% of PBS treated mice succumbed to LPS after 24 hours while NAD^+^ treated animals exhibited an overall survival >85% (**Fig. 1D**), which was consistent with our bacterial infection model. Mice infected and treated with NAD^+^ survived for several months and recovered fully after 10 days. Of note, mice survived for over a year following infection and died of aging.

**Figure 1.**
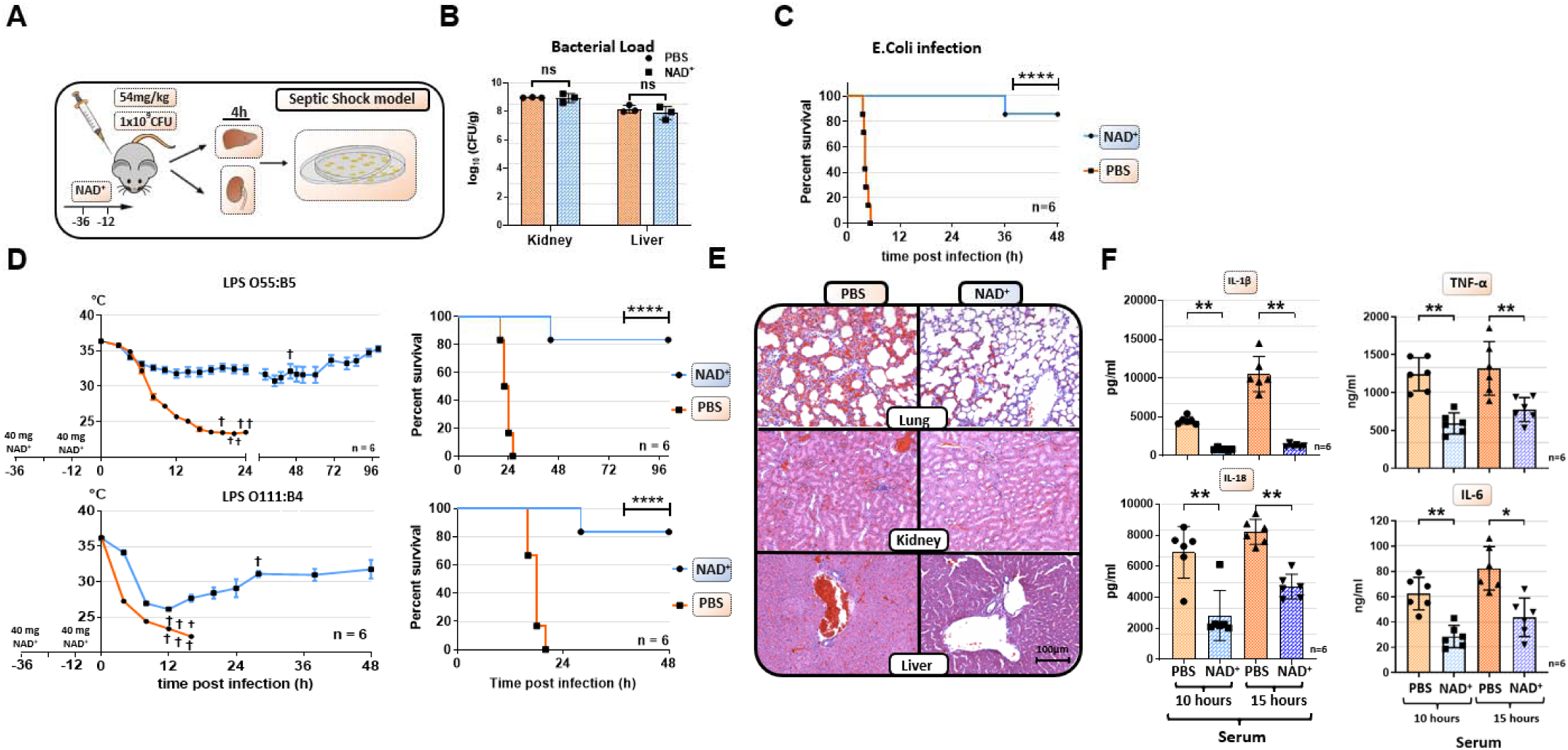
NAD^+^ protects mice from lethal bacterial infection and endotoxic shock by dampening systemic inflammation. **(A)** mice were treated with PBS or NAD^+^ prior to administration of a lethal dose of either pathogenic *E. coli* or LPS by intraperitoneal injection. **(B)** Kidneys and Livers were removed after and bacterial load was determined by counting CFU **(C)** Survival was monitored over 48 hours after bacterial infection and **(D)** LPS injection of both serotypes (n=6, 3 independent survival experiments). In addition, body temperature was monitored in the kinetics of up to 100 hours **(E)** Lungs, Kidneys and Livers were removed IHC was performed for H&E staining **(F)** Systemic levels (serum) of IL-6, TNFα, IL-1β and IL-18 were assessed by ELISA. Column plots display mean with standard deviation (n=5). Statistical significance was determined by using Student’s T-test or One-Way-ANOVA while survival data were compared using log-rank Mantel-Cox test. Asterisks indicate p-values * = p<0.05, **= p<0.01 and *** = p<0.001, only significant values are shown. All data depicted in this figure are provided as source data. Figure 1 Source Data 1 – Raw Data for **Fig. 1B**: Bacterial Load Figure 1 Source Data 2 – Raw Data for **Fig. 1C**: E. coli Infection Figure 1 Source Data 3 – Raw Data for **Fig. 1D**: LPS Infection Figure 1 Source Data 4 – Raw Data for **Fig. 1E**: Histology Figure 1 Source Data 5 – Raw Data for **Fig. 1F**: ELISA

LPS-induced death derives from multi-organ failure^20^. Therefore, lung, kidney, liver, ileum and spleen were harvested 15 hours after LPS administration and tissue damage was assessed by Hematoxylin and Eosin (H&E) staining. Tissue evaluation indicated severe pulmonary hemorrhage, excessive tubular fibrin deposition, hepatocyte cell swelling, disseminated intravascular coagulation (DIC) and ileal villi destruction consistent with a multi-organ-dysfunction syndrome (MODS)^21^ in mice treated with PBS. In contrast, NAD^+^ administration dramatically attenuated signs of organ failure with significantly less pulmonary hemorrhage and DIC, intact liver and kidney tissue architecture and preserved ileal villi (**Fig. 1E**, **Fig. 1 Supplement 1 and Fig. 1 Supplement 2**).

To elucidate the protective effects of NAD^+^ systemic levels of IL-1β and IL-18, two major cytokines implicated in inflammasome activation, were measured 10 and 15 hours after intraperitoneal injection of LPS (**Fig. 1F**). Of, note IL-6 and TNFα systemic levels were measured as well (**Fig. 1F**).

Our findings indicated that LPS injection resulted in a robust systemic increase of IL-1β, IL-6, TNFα and IL-18 in the PBS group, which was almost abolished in NAD^+^ treated mice. Taken together, our results suggest that NAD^+^ protects mice against septic shock not via bacterial clearance but rather via inflammasome blockade.

### NAD^+^ specifically inhibits the non-canonical inflammasome

Our data suggest that NAD^+^ protects against septic shock via inflammasome blockade. Monocytes, especially macrophages, have been described as major drivers of inflammasome derived cytokine secretion in the context of septic shock^22^. Thus, to test the effect of NAD^+^ on inflammasome function, bone marrow derived macrophages (BMDMs) were obtained and both, canonical and non-canonical inflammasomes were stimulated in in the presence or absence of 100 µmol/ml NAD^+^. Activation of the canonical pathway was achieved through LPS priming (1µg/ml) followed by ATP stimulation (5 mmol/l). Notably, BMDMs subjected to NAD^+^ or PBS treatment followed by canonical inflammasome activation did not exhibit any significant difference in IL-1β secretion or pyroptosis that was assessed by LDH release measurement, a marker for cell death^23^ (**Fig. 2A**).

**Figure 2.**
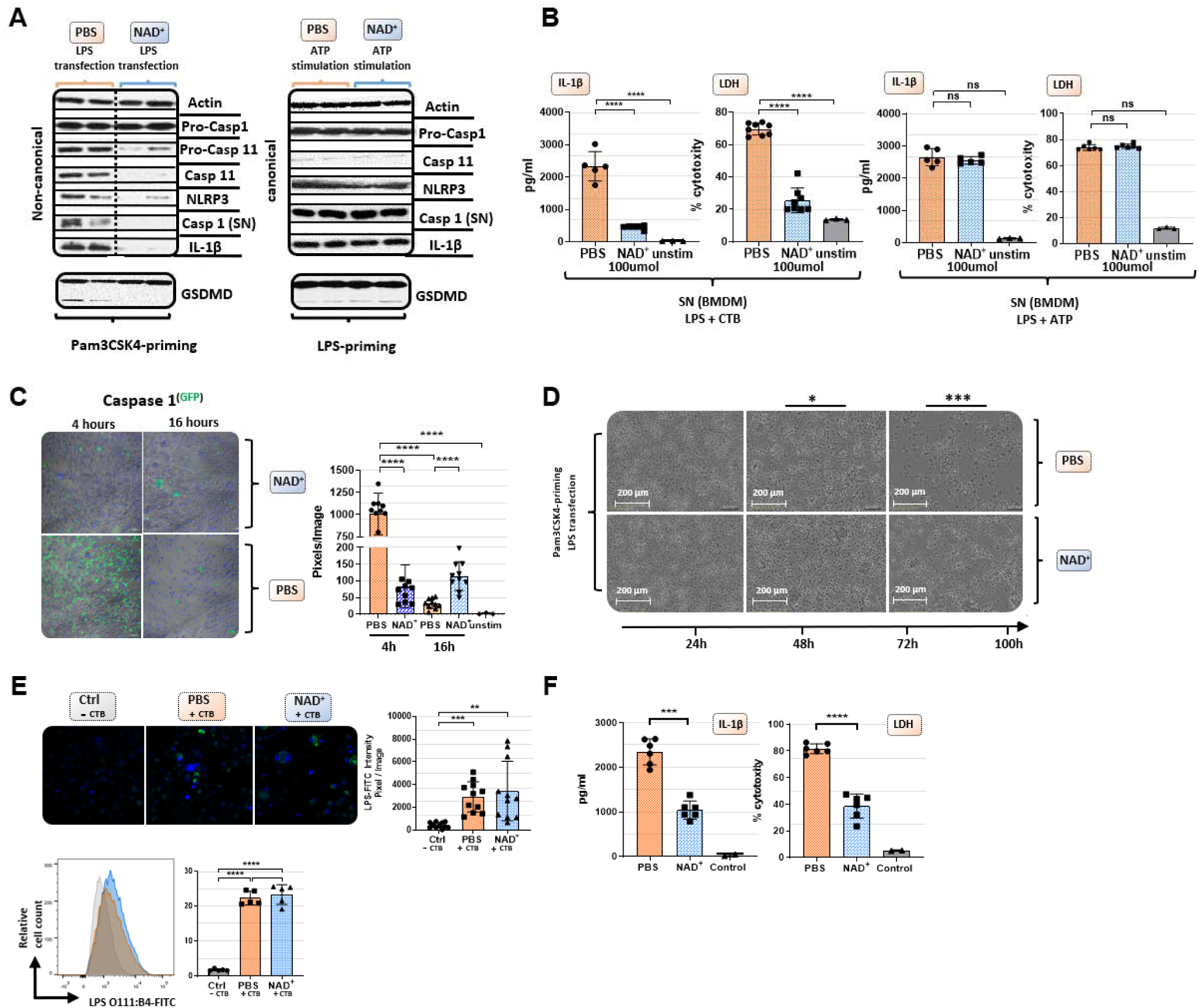
NAD^+^ specifically inhibits the non-canonical inflammasome by targeting caspase 11. Bone marrow was isolated from mice and BMDM were differentiated in vitro. Subsequently, BMDM were cultured in the presence of NAD^+^ or PBS. BMDMs were then primed with either Pam3CSK4 or LPS O111:B4. Next primed BMDMs were stimulated with ATP or LPS and CTB. **(A)** Pro-casp-1, Pro-casp-11, Casp-11, NLRP3, Casp-1, IL1β and GSDMD expression were determined using Western blot and **(B)** IL-1β secretion and LDH release were assessed in the supernatant. **(C)** Time dependent Caspase-1 expression was determined via active staining and assessed using a confocal microscope. **(D)** Cell viability and apoptosis were monitored using the IncuCyte^®^ live microscopy system. **(E)** LPS transfection ith CTB was visualized by using FITC-coupled LPS and DAPI staining and quantified by confocal microscopy and flow cytometry. **(F)** For human experiments macrophages were differentiated from PBMC, primed with Pam3CSK4 and subsequently transfected with LPS and 0.25% Fugene HD Plus. Column plots display mean with standard deviation (n=6). Statistical significance was determined by using Student’s T-test or One-Way-ANOVA. Asterisks indicate p-values * = p<0.05, **= p<0.01 and ***= p<0.001, only significant values are shown. All data depicted in this figure are provided as source data. Figure 2 Source Data 1 – Raw Data for **Fig. 2A**: Original Western Blots Figure 2 Source Data 2 – Raw Data for **Fig. 2A**: Western Blots with highlighted Bands and Sample Labels Figure 2 Source Data 3 – Raw Data for **Fig. 2B**: ELISA Mouse BMDMs Figure 2 Source Data 4 – Raw Data for **Fig. 2C**: Caspase 1 Staining Figure 2 Source Data 5 – Raw Data for **Fig. 2D**: Incucyte Live Microscopy Figure 2 Source Data 6 – Raw Data for **Fig. 2E**: LPS Transfection Staining Figure 2 Source Data 7 – Raw Data for **Fig. 2F**: ELISA Human Macrophages

To trigger the non-canonical inflammasome pathway, BMDMs were primed with Pam3CSK4, a TLR1/2 agonist, followed by cholera toxin B (CTB) and LPS (2µg/ml) administration. The data showed that NAD^+^ treatment resulted in a robust reduction of IL-1β release and cell death when compared to the PBS control group (**Fig. 2A**). Furthermore, Western blotting revealed that BMDMs cultured in presence of NAD^+^ exhibited a dramatic decrease of casp-11 expression and its downstream targets including casp-1, IL-1β and cleaved gasdermin-D (GSDMD) (**Fig 2B**).

Moreover, we observed a prominent decrease in casp-1 expression under NAD^+^ treatment that was constant over the time course of 16 hours. In contrast, BMDMs treated with PBS exhibited excessive casp-1 expression at 4 hours that was found to be absent after 16 hours (**Fig. 2C**), which is consistent with the strong cytotoxicity leading to membrane permeabilization and release of Casp-1 into the supernatant. Noteworthy, Pam3CK4 derived BMDM priming was not affected by NAD^+^ since NF-κB as well as pro-caspase-1 levels had not been altered (**Fig. 2A and Fig. 2 Supplement 1**) underlining the specific inhibition of casp-11. To visualize NAD^+^ mediated blockade of pyroptotic macrophage death, BMDMs were treated with PBS or NAD^+^, primed with Pam3CSK4, then stimulated with LPS and CTB and cell viability and apoptosis were monitored using the IncuCyte^®^ live microscopy system. Hereby, we observed distinct longitudinal kinetics over 100 hours with complete disaggregation of cell integrity in the PBS group contrary to overall preserved cell structure in NAD^+^ treated BMDMs (**Fig. 2D**, **Fig. 2 Supplement 2, Mov. 1**). To rule out, that NAD^+^ impairs LPS internalization into cells, BMDMs were stimulated with CTB and LPS that was coupled to a fluorescent reporter (FITC) and transfection effectivity was assessed by fluorescence microscopy and flow cytometry.

Our data indicated no significant difference between the PBS and NAD^+^ treated group (**Fig. 2E**), suggesting that NAD^+^ does not alter LPS internalization. Notably, BMDMs only stimulated with LPS showed no internalization of LPS consistent with previous reports^12^.

Casp-4 and 5 have been delineated as the human homolog of casp-11 in mice carrying out the same effector functions including pyroptosis induction and IL-1β secretion^24^. As clinical relevance, we therefore tested whether NAD^+^ was also able to block the non-canonical pathway in human macrophages. Hence, human macrophages were differentiated from PBMC and treated with NAD^+^ followed by intracellular LPS transfection (Fugene) and IL-1β secretion and cytotoxicity were quantified. The results indicated that NAD^+^ treatment significantly dampened both, IL-1β secretion and pyroptosis (**Fig. 2F**), underscoring its therapeutic potential. Collectively, our results suggest that NAD^+^ acts directly on macrophages by targeting specifically the non-canonical inflammasome signaling machinery.

### NAD^+^ inhibits the non-canonical inflammasome via STAT-1/IFN-***β*** pathway blockade

Although our data emphasized that NAD^+^ blocks the non-canonical inflammasome pathway, the underlying mechanisms remained yet to be determined. Therefore, we performed RNA-sequencing of Pam3CSK4 primed BMDMs that were treated with PBS or NAD^+^ and subsequently stimulated with CTB + LPS O111:B4. Interestingly, when blotting gene expression differences in a Venn diagram, we found strikingly more genes commonly expressed in the NAD^+^ and control group when compared to the PBS treated group (**Fig. 3A**). Gene ontology enrichment analysis revealed a significant downregulation of genes involved in the antiviral response in addition to the cellular response to the type-I-IFN, IFN-β, when comparing NAD^+^ and PBS treated groups (**Fig. 3B**). Type-I-IFN are known to promote the expression of over 2,000 IFN-stimulated genes (ISGs), translated into ISGs-induced proteins which have been shown to act by enhancing pathogen detection and restrict their replication^25^. Recently, it was reported that type-I-IFNs are required for casp-11 expression contributing to non-canonical inflammasome activation^26, 27^. Consistently, LPS-stimulated macrophages from TRIF-deficient mice displayed impaired casp-11 expression, implying a context-dependent role for type-I-IFN in the regulation of caspase-11 activity^26^. Indeed, when comparing expression of genes involved in IFN-β signaling through cluster analysis we found a significant decrease in a broad range in genes in the NAD^+^ treated group (**Fig. 3C**). Most strikingly, GTPases and guanylate binding proteins involved in the downstream signaling of IFN-β were significantly downregulated (**Fig. 3C and Fig. 3D**) while IFN-β-receptor 2 expression remained unaffected (**Fig. 3C**). Recently, IFN-inducible GTPases and guanylate binding proteins have been assigned a crucial role for the intracellular recognition of LPS and linked caspase-11 activation^27, 28^. Thus, to test if NAD^+^ mediated non-canonical inflammasome blockade via IFN-β, NAD^+^ or PBS treated BMDMs were primed with Pam3CSK4 and subsequently stimulated with LPS O111:B4 + CTB and 1000 U/ml of recombinant IFN-β. Strikingly, administration of recombinant IFN-β resulted in a complete reversal of NAD^+^ mediated blockade of IL-1β secretion and pyroptosis (**Fig. 3E**). Moreover, IFN-β administration restored casp-11, NLRP3 and GSDMD expression in the NAD^+^ treated group (**Fig. 3F**). It is well established that STAT-1 phosphorylation constitutes the link between intracellular type-I-IFN signaling and the transcription of ISGs through nuclear translocation^29, 30^. Notably, our RNA-seq data indicated a significant downregulation of STAT-1 (**Fig. 3C**).

**Figure 3.**
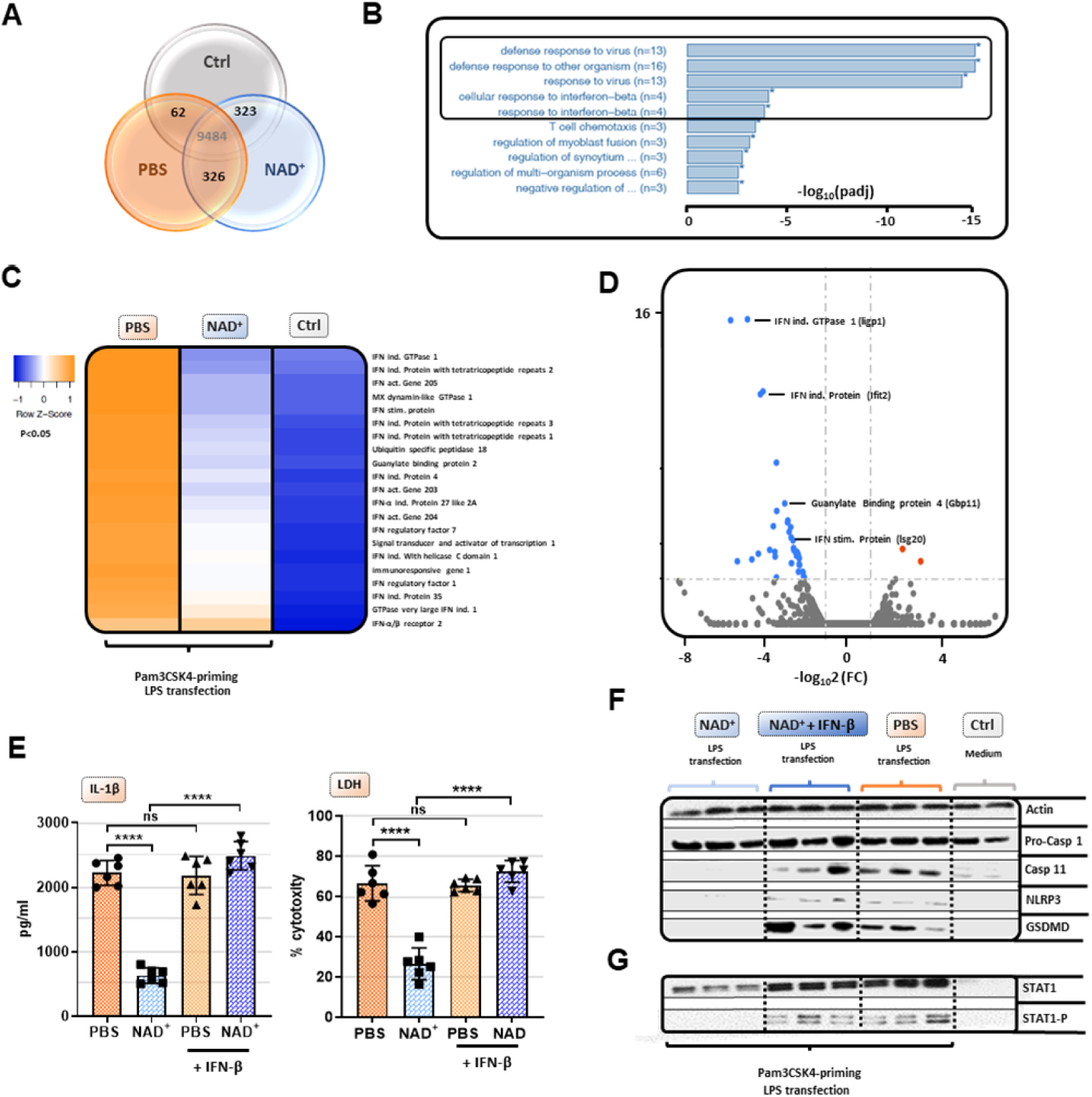
NAD^+^ mediated inhibition of the non-canonical inflammasome is based on an impaired response to IFN-β. Differentiated BMDMs were cultured in the presence of NAD^+^ or PBS. BMDMs were then primed with Pam3CSK, subsequently stimulated with LPS and CTB and RNA sequencing was performed. Unstimulated BMDMs served as controls. **(A)** Venn diagram plotting common gene expression between all 3 groups **(B)** Gene ontology enrichment analysis displaying the highest significant pathways differing when comparing NAD^+^ and PBS treated BMDMs **(C)** Expression cluster analysis of genes involved in IFN-β signaling through cluster analysis depicted in a heat map **(D)** Vulcano plot displaying the most significant genes up or downregulated comparing NAD^+^ and PBS treated BMDMs **(E)** Stimulated BMDMs were additionally treated with recombinant INF-β, and IL-1β and LDH release were measured. **(F)** Moreover, pro-Casp-1, Casp-11, NLRP3, GSDMD, **(G)** STAT-1 and Phospho-STAT-1 expression were assessed by Western blot. Column plots display mean with standard deviation (n=6). Statistical significance was determined by using Student’s T-test or One-Way-ANOVA. Asterisks indicate p-values * = p<0.05, **= p<0.01 and ***= p<0.001, only significant values are shown. All data depicted in this figure are provided as source data. Figure 3 Source Data 1 – Raw Data for **Fig. 3E**: ELISA BMDM Figure 3 Source Data 2 – Raw Data for **Fig. 3F**: Original Western Blots Figure 3 Source Data 3 – Raw Data for **Fig. 3F**: Western Blots Highlighted Bands and Sample Labels Figure 3 Source Data 4 – Raw Data for **Fig. 3G**: Original Western Blots Figure 3 Source Data 5 – Raw Data for **Fig. 3G**: Western Blots Highlighted Bands and Sample Labels

Moreover, we have previously shown that NAD^+^ administration dampens the expression and activation of transcription factors such as STAT-5^16^. To test, whether NAD^+^ blocks IFN-β signaling via STAT-1, BMDMs were subjected to NAD^+^ or PBS followed by non-canonical inflammasome stimulation and recombinant IFN-β. After 16 hours STAT-1 expression and phosphorylation were assessed by Western blotting. Consistent with our previous results, NAD^+^ treatment downregulated expression levels of STAT-1 and phospho-STAT-1. In contrast, addition of recombinant IFN-β treatment to NAD^+^ treated BMDMs restored STAT-1 and phospho-STAT-1 expression that was equivalent to the PBS treated group (**Fig. 3G**). Taken together, our data indicate that NAD^+^ impedes non-canonical inflammasome activation via IFN-β /STAT-1 blockade (**Fig. 4)**.

**Figure 4.**
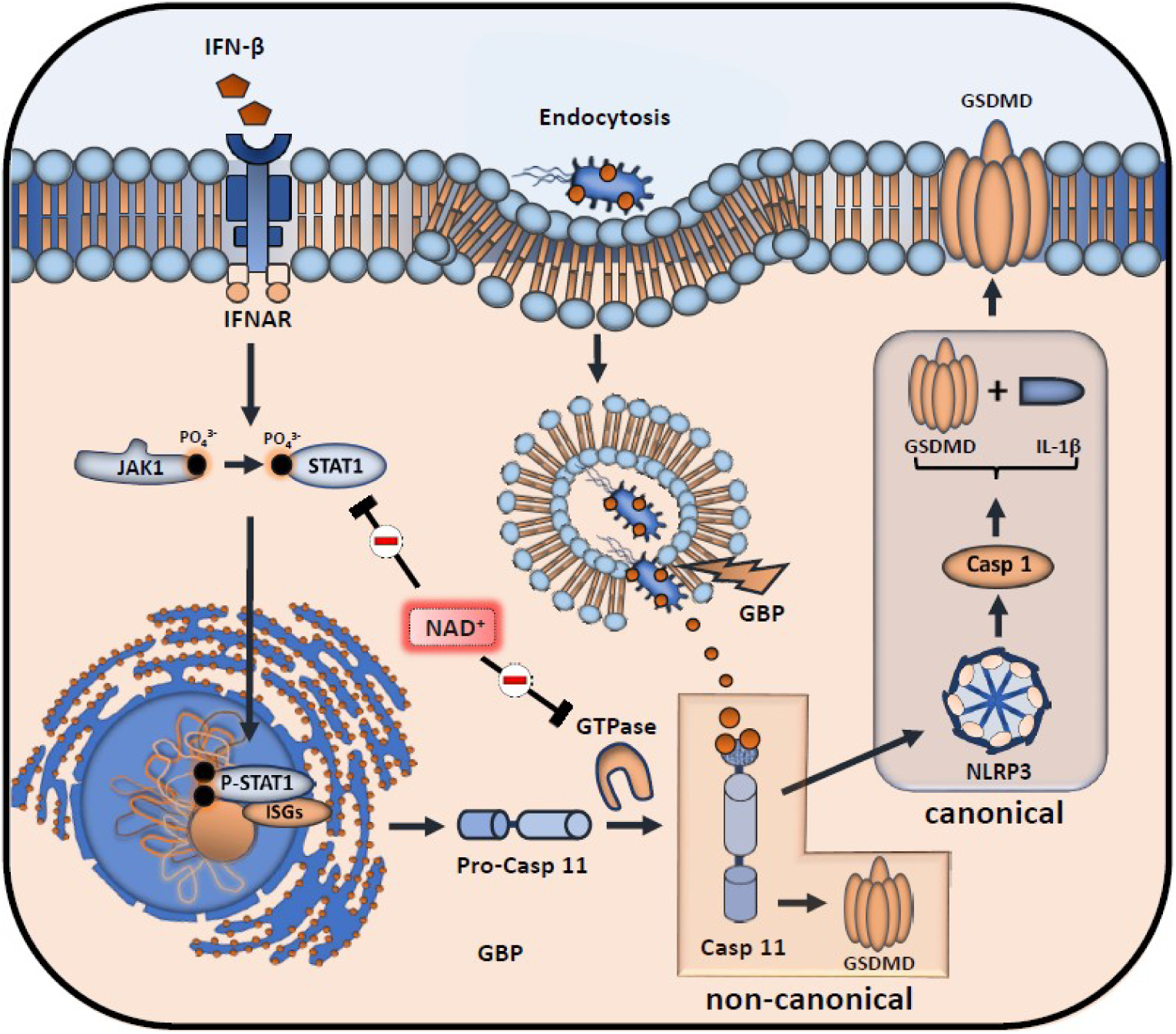
Inhibitory effects of NAD^+^ on IFN-β downstream signaling and inflammasome activation. NAD^+^ inhibits STAT-1 expression and phosphorylation, thus compromising the intracellular response to IFN-β. Subsequently, stimulation of the IFNAR receptor by IFN-β leads to a decreased transcription of pro-caspase-11 as well as ISGs (IFN-inducible GTPases and GBPs). Due to diminished caspase-11 levels, non-canonical inflammasome activation through intracellular, gram-negative bacteria opsonization by GBPs is significantly inhibited.

### NAD^+^ increases Caspase-1 KO mice resistance to endotoxic shock via systemic IL-10 production

Caspase-11 KO mice have been reported to be resistant towards lethal doses of LPS inducing septic shock^12^. However, upon priming with TLR3 instead of a TLR4 ligand, Casp-11 KO mice merely exhibit partial resistance towards LPS-induced shock with a 50–60% survival rate^12, 31^. Our data indicate that NAD^+^ prevents LPS-induced cell death via the non-canonical inflammasome pathway and casp-11 blockade. We thus tested whether NAD^+^ could achieve similar protection against septic shock in WT vs casp-11 KO mice. Casp-11 KO mice were intraperitoneally injected with NAD^+^ and PBS and treated with 6 mg/kg poly(I:C) 6 hours prior LPS administration.

Consistent with previous studies the results indicated a modest resistance of casp-11 KO mice (40% survival). In high contrast, both WT and casp-11 KO mice subjected to NAD^+^ exhibited 85%-100% survival, respectively when compared to casp-11 KO mice that were treated with PBS, suggesting the existence of an alternative protective pathway against septic shock that is casp-11 independent. WT mice, treated with 6 mg/kg poly(I:C) followed by LPS (54mg/kg) administration not only survived but fully recovered 7 days later, underscoring the unique and robust therapeutic effect of NAD^+^ in septic shock.

Previous studies have reported inferior outcomes of IL-10 ^-/-^ mice in septic shock^32, 33^ pointing out a 20-fold lower lethal dose of LPS compared to WT mice^33^. Moreover, IL-10 itself has been shown to prevent mice from septic shock induced death after a single administration^34^. We have previously delineated immunosuppressive properties of NAD^+^ via a systemic production of IL-10, a robust immunosuppressive cytokine. In addition, we have described the pivotal role of NAD^+^ protecting towards EAE and allograft rejection via an increased frequency of IL-10 producing CD4^+^ T cells^15, 16^. To test if IL-10 plays an additional protective role in the context of NAD^+^ mediated protection towards LPS-induced death, WT mice treated with NAD^+^ or PBS were subjected to intraperitoneal LPS injection (54mg/kg) and IL-10 expression by macrophages, dendritic cells and T cells was assessed 15 hours after LPS administration.

Consistent with our previous studies^15, 16^, we found significantly augmented frequencies of IL-10 producing CD4^+^ and CD8^+^ T cells (**Fig. 5C**). Moreover, we detected a dramatic increase of IL-10 production by macrophages, but not the DC population (**Fig. 5B**). Interestingly, IL-10 has been described to inhibit macrophage function and pro-inflammatory cytokine production in both, human^35^ and mice^36^. Moreover, autocrine IL-10 secretion of macrophages was found to decrease pro-IL-1β concentration by promoting signal transducer activator of transcription-3 (STAT-3) expression^37^. To investigate the potential autocrine impact of an augmented IL-10 production on macrophage self-regulation, we administered combined IL-10 neutralizing antibody and IL-10 receptor antagonist to BMDMs primed with Pam3CSK4 and stimulated with CTB and LPS O111:B4. The results showed that neutralization of the autocrine IL-10 signaling pathway dampened NAD^+^ mediated decrease of IL-1β secretion and reversed pyroptotic cell death partially (**Fig. 5D**). To further investigate the relevance of our in vitro findings, IL-10^-/-^ mice were treated with NAD^+^ or PBS, subjected to LPS (54mg/kg) and survival was monitored. Consistent with previous reports^32, 33^, mice lacking IL-10 exhibited an inferior protection against septic shock when compared to WT animals. More importantly, IL-10^-/-^ mice subjected to NAD^+^ exhibited a compromised survival (**Fig. 5E**) suggesting that systemic production of IL-10 following NAD^+^ administration plays a pivotal role in NAD^+^-mediated protection against septic shock.

**Figure 5.**
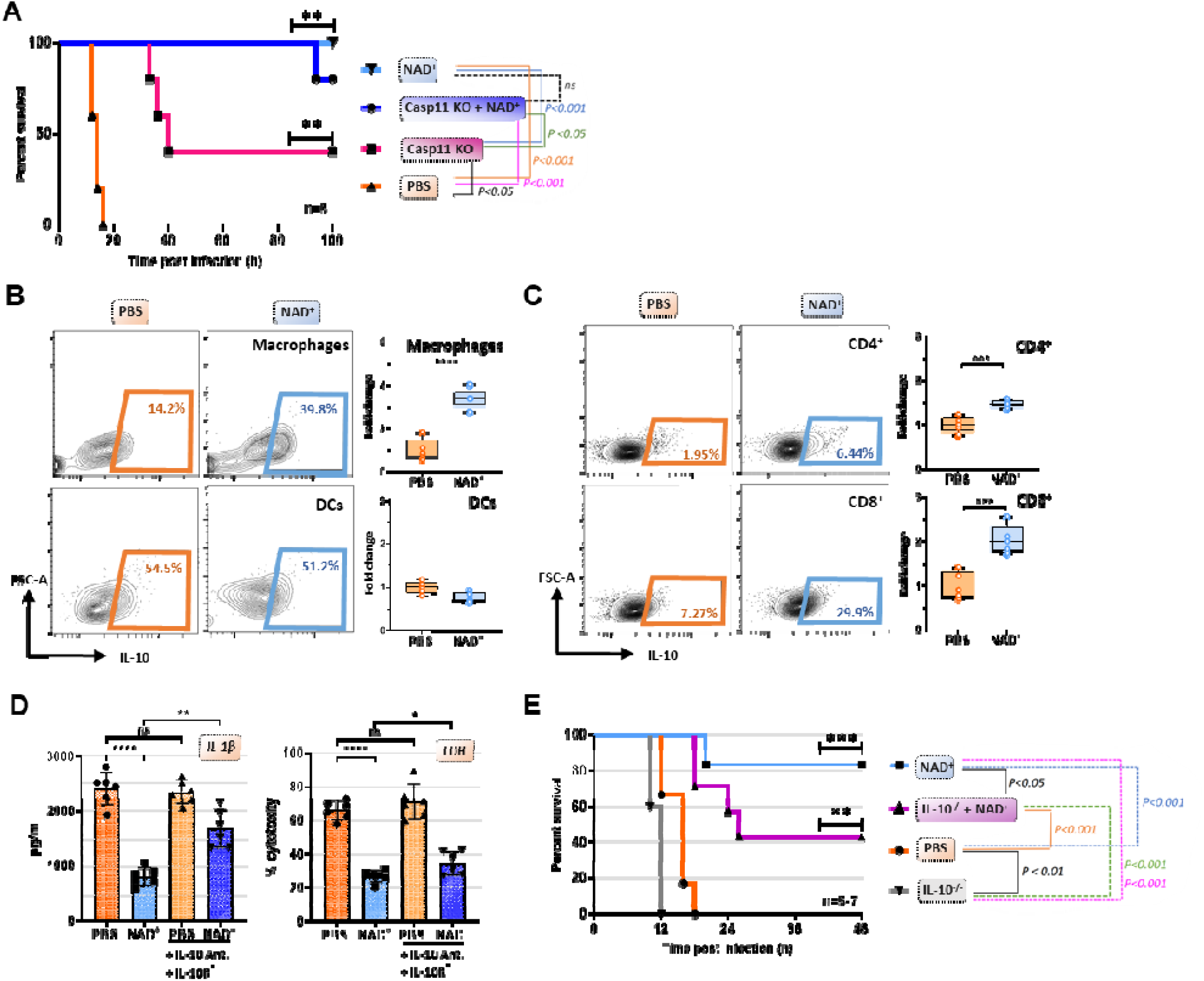
IL-10 constitutes an additional mechanism mediating the protective capacities of NAD^+^ in the context of septic shock. **(A)** Caspase-11 KO (knockout) mice were treated with NAD^+^ or PBS. Subsequently mice were subjected to Poly(I:C) prior to LPS injection and survival was monitored (n=5, 2 independent survival experiments). Mice treated with either NAD^+^ or PBS were injected with LPS and after 10 hours, Splenic frequencies of IL-10 producing **(B)** Macrophages and Dendritic cells **(C)** and CD4^+^ and CD8^+^ T cells were assessed by flow cytometry. **(D)** BMDMs treated with NAD^+^ or PBS were stimulated with LPS and CTB in the presence of IL-10 neutralizing antibodies and IL-10 receptor antagonists. Subsequently IL-1β and LDH release were assessed. **(E)** IL-10^-/-^ mice treated with NAD^+^ or PBS were challenged with LPS and survival was monitored (n=5-7, 2 independent survival experiments). Column plots display mean with standard deviation (n=6). Statistical significance was determined by using Student’s T-test or One-Way-ANOVA while survival data were compared using log-rank Mantel-Cox test. Asterisks indicate p-values * = p<0.05, **= p<0.01 and *** = p<0.001, only significant values are shown. All data depicted in this figure are provided as source data. Figure 5 Source Data 1 – Raw Data for **Fig. 5A**: Casp11 KO Survival Figure 5 Source Data 2 – Raw Data for **Fig. 5B**: FACS Macrophages and DCs Figure 5 Source Data 3 – Raw Data for **Fig. 5C**: FACS CD4+ and CD8+ T cells Figure 5 Source Data 4 – Raw Data for **Fig. 5D**: ELISA BMDM Figure 5 Source Data 5 – Raw Data for **Fig. 5E**: IL-10^-/-^ Survival

## Discussion

Previously, we have delineated the protective role of NAD^+^ in the context of *L. m.* infection, a gram-positive bacterium^17^. However, it remained unclear whether NAD^+^ conveyed resistance towards *L. m.* by an augmented bacterial clearance or rather through its immunosuppressive effects dampening pathological systemic inflammation. Although the cell membrane of *L. m.* has been shown to bear lipoteichoic acids, which resemble the endotoxin LPS from gram-negative bacteria in both, structure and function, it is widely considered as an intracellular bacterium^38^. In our current study, we administered a lethal dose of pathogenic *E. coli*, that is well known to promote septic shock, and showed that NAD^+^ also protected towards a lethal dose of this gram-negative bacterium. More importantly, we demonstrate that NAD^+^ conveys protection towards septic shock by specifically inhibiting the non-canonical inflammasome but not via bacterial clearance. Mechanistically, NAD^+^ impedes pro-casp-11 and casp-11 expression in macrophages blocking non-canonical derived GSDMD cleavage and NLRP3 inflammasome activation, thus inhibiting pyroptotic cell death and pro-inflammatory cytokine release. The resistance of NAD^+^ treated WT mice against *E. coli* and LPS induced septic shock reflected the robust inhibitory effect observed in vitro of NAD^+^ on the non-canonical inflammasome signaling machinery.

Until now, the exact mechanism how pro-casp-11 expression and its maturation to casp-11 is regulated remains unclear. Given the low basal expression of both pro-casp-11 and casp-11^39^, a priming signal is required for initiating the non-canonical inflammasome pathway and macrophage sensing of intracellular LPS^40^. Previous work has demonstrated that transcriptional induction of the pro-casp-11 isoforms p42 and p38 in macrophages requires type I IFN stimulation^39, 41^ while IFN-β has been shown to promote transcriptional induction and processing of caspase-11^26^. In line with these findings, CTB treatment of macrophages primed with Pam3csk4 failed to elicit IL1-β release compared to LPS primed controls while exogenous administration of IFN-β in turn restored CTB-induced IL-1β production^26^ underscoring the transcriptional role of type I IFN. Our RNA-seq results indicated a dampened cellular response towards IFN-β while western blotting revealed a significant downregulation of both, pro-casp-11 and casp-11 suggesting a transcriptional downregulation of both enzymes. Consistently, NAD^+^ decreased STAT-1 expression and phosphorylation, which constitutes the mechanistic link between extracellular type I IFN stimulation and transcriptional effects through translocation of phosphorylated STAT-1 to the nucleus inducing ISGs^30^. Thus, treatment of stimulated macrophages subjected to NAD^+^ with recombinant IFN-β restored STAT-1 signaling, caspase-11 expression and GSDM cleavage which translated into reconstituted IL-1β production and LDH release. Collectively, NAD^+^ mitigates the intracellular response to IFN-β that leads to non-canonical inflammasome induction by suppressing macrophage derived STAT-1 expression and phosphorylation.

Furthermore, we showed that NAD^+^ treatment improved resistance of casp-11 KO mice towards poly I:C primed septic shock. More importantly, WT mice treated with NAD^+^ exhibited 100% survival while casp-11 KO mice treated with PBS exhibited a modest 40% survival, suggesting that NAD^+^ promotes survival beyond non-canonical inflammasome blockade. Our previous studies have delineated the effects of NAD^+^ on various immune cells such as dendritic cells and CD4^+^ T cells including Th1, Th17, regulatory type 1 (Tr1) and Treg cells communicated exclusively through mast cells (MCs)^15, 16, 17^. Thereby, NAD^+^ treatment promoted MC derived induction of TR1 cells that resulted into increased systemic levels of IL-10. Latter one was found to play a cardinal feature during bacterial infection as MC^-/-^ mice were more susceptible to *L. m.* infection than WT animals when treated with NAD^+^. Here, we found a direct effect of NAD^+^ on macrophages by specifically inhibiting the non-canonical inflammasome and promoting IL-10 production. Polymorphisms in the IL-10 locus or IL-10R deficiencies have been linked to severe intestinal inflammatory diseases in both, human and mice^42, 43, 44, 45^. More importantly, mice deficient for IL-10 have been shown to display elevated inflammasome activation and IL-1β production resulting in severe colitis^46^ or Ag-induced arthritis^47^. When inhibiting the autocrine pathway for IL-10 through combined receptor antagonization and IL-10 neutralization, we found a pronounced increase of IL-1β production of NAD^+^ treated macrophages stimulated with CTB and LPS (Fig. 4D). This is consistent with previous reports showing that autocrine IL-10 signaling interferes with the transcription of pro-IL-1β^37^. LDH release in turn, was only restored partly possibly due to missing effects of second party leucocytes secreting IL-10 in vivo such as Tr1 cells which have been shown to inhibit the transcription of IL-1β and inflammasome mediated activation of caspase-1^48^.

More recently, casp-8, that plays a central role in apoptosis, has been reported as an important mediator of endotoxemia resistance and LPS-driven systemic inflammation. Since our RNA-sequencing results revealed a dramatically attenuated cellular response towards type-I-IFN with downregulation of various interferon regulatory factors, that have been reported as major regulators of casp-8^49, 50^ it is possible that NAD^+^ may exert protection against septic shock by altering caspase-8 expression as well. Although, we have previously reported the protective effect of NAD^+^ against apoptosis of activated CD4^+^ T cells^15^, it remains yet to be determined how NAD^+^ impacts executioner proteins of other cell death processes such as apoptosis and necroptosis.

Notably, both casp-8 and casp-11 have been found dispensable in the hematopoietic compartment that produces the pro-inflammatory cytokines necessary to initiate shock^51^. Thus, NAD^+^ treatment may improve resistance of casp-11 KO mice to septic shock by also dampening the initiating pro-inflammatory cytokine cascade via its systemic IL-10 cytokine production.

Importantly, while inhibiting macrophage derived inflammasome function, NAD^+^ does not interfere with NF-κB signaling which has been shown to promote various inflammatory and autoimmune diseases when dysregulated^52^.

Taken together, we dissected the dichotomous capacity of NAD^+^ to dampen auto-and allo-immunity while concomitantly protecting towards severe bacterial infection, outlining its unique effects in the context of septic shock.

## Material and Methods

### Animals

Young (8-12 weeks) C57BL/6, B6.129P2-IL10tm1Cgn/J and B6.129S4(D2)-Casp4tm1Yuan/J mice were purchased from Charles River Laboratory, Wilmington, MA. All mice were male, age-matched and experimental and control animals were housed separately. Animals and samples were randomly assigned to either the control or treatment group to ensure biological diversity. The study protocol was approved by the Brigham and Womeńs Hospital Institutional Animal Care and use Committee (IACUC) (animal protocol #2018N000049). All mice were male, age-matched and experimental and control animals were housed separately. Owing to the exploratory nature of our study, we did not use randomization and blinding. No statistical methods were used to predetermine sample size. All animals were maintained in specific pathogen-free (SPF) conditions at the Brigham and Womeńs Hospital animal facility in accordance with federal, state, and institutional guidelines. Animals were maintained on 12-h light, 12-h dark cycle in facilities with an ambient temperature of 19–22 °C and 40–60% humidity and were allowed free access to water and standard chow. Euthanasia was performed by cervical dislocation following anesthesia with isoflurane (Patterson Veterinary, Devens, MA, USA).

### Murine Bone Marrow Derived Macrophage Differentiation and Culture

8-12-week-old C57BL/6 mice were euthanized by cervical dislocation, sprayed with alcohol and skin was removed to expose femurs. The femur was flushed with ice cold PBS and the obtained bone marrow was filtered through 70um Nylon cell strainer. After washing with PBS, red blood cell lysis was performed using ammonium-chloride-potassium-solution (Fisher scientific) and the reaction was blocked with complete Dulbecco’s modified eagle medium (DMEM) (Fisher Scientific) supplemented with 10% endotoxin-free bovine serum and PS. To minimize fibroblast contamination cells were cultured in complete DMEM at 37°C, 5% CO_2_ and non-adherent cells were collected after 30 minutes.

Bone marrow cells were then differentiated into macrophages in DMEM supplemented with 10% endotoxin-free bovine serum, PS and 40ng/ml murine GM-CSF (Abcam) for 8 days. Medium was changed every 2 days to remove non-adherent cells.

### Canonical and Non-canonical Inflammasome Activation in Murine Macrophages

After 8 days of culture the medium was replaced by 40ng/ml GM-CSF containing 100µmol NAD^+^ culture medium. For 2 following days NAD^+^ was added daily until stimulation. To induce canonical and non-canonical inflammasome activation in murine macrophages, NAD^+^ treated and control BMDMs were cultured overnight in a 24 well plate at 1×10^6^ cells/ml and afterwards primed with 1µg/ml Pam3CSK4 or 1µg/ml LPS O111:B4 (Sigma) for 5-6 hours. Primed BMDMs were then stimulated for 16 hours with either 5 mmol ATP (canonical inflammasome stimulation) or 2µg/ml LPS O111:B4 and 20µg/ml CTB (Sigma) to allow LPS entry (non-canonical inflammasome stimulation) where indicated. To test the effect of NAD^+^ on type 1 IFN and STAT1 signaling, BMDMs were cultured overnight in a 24 well plate at 1×10^6^ cells/ml and afterwards primed with 1µg/ml Pam3CSK4 or 1µg/ml LPS O111:B4 (Sigma) for 5-6 hours. Primed BMDMs were then stimulated for 16 hours 2µg/ml LPS O111:B4, 20µg/ml CTB and U/ml recombinant IFN-β.

### ELISA

Expression of macrophage derived murine IL-1β, IL-18 and human IL-1β was analyzed in the cell culture supernatant by commercial ELISA kits (Invitrogen) following the manufacturer’s recommended procedures.

### Pyroptosis Assay

Pyroptotic cell death was measured by assessing LDH-release in the cell culture supernatant of human and murine macrophages using a CytoTox 96 Non-radioactive Cytotoxic Assay (Promega) following the manufacturer’s recommended procedures.

### RNA Extraction and RNA-Sequencing

BMDMs were harvested and differentiated as outlined in the particular section. After 8 days of culture the medium was replaced by 40ng/ml GM-CSF containing culture medium (control group) or 40ng/ml GM-CSF containing 100µmol NAD^+^ culture medium (NAD^+^ treated group). For 2 following days NAD^+^ was added daily. Subsequently, NAD^+^ treated and control BMDMs were cultured overnight in a 24 well plate at 1×10^6^ cells/ml and afterwards primed with 1µg/ml Pam3CSK4 or 1µg/ml LPS O111:B4 (Sigma) for 5-6 hours. Primed BMDMs were then stimulated for 16 hours with 2µg/ml LPS O111:B4 and 20µg/ml CTB (Sigma) to allow LPS entry. Another set of BMDMs were differentiated without additional treatment serving as naive controls.

Subsequently, RNA was extracted with the RNAqueous extraction kit (Applied Biosystems), according to the manufacturer’s protocols. Briefly, cells were homogenized in lysis buffer (total volume of 0.5 mL) and passed through a column. After successive washes, RNA was eluted. RNA sequencing was commercially performed by Novogene CO., Ltd. In brief, mRNA was enriched from total RNA using oligo(dT) beads and subsequently fragmented randomly in fragmentation buffer, followed by cDNA synthesis using random hexamers and reverse transcriptase. After first-strand synthesis, a custom second-strand synthesis buffer (Illumina) was added with dNTPs, RNase H and Escherichia coli polymerase I to generate the second strand by nick-translation. The final cDNA library is ready after a round of purification, terminal repair, A-tailing, ligation of sequencing adapters, size selection and PCR enrichment.

### Isolation and Differentiation of Human Macrophages from PBMCs

Blood was obtained from healthy adult volunteers with the only purpose to isolate PBMCs in order to create a basis for macrophage cultures. Blood withdrawal was performed in accordance with guidelines of and approved by the Institutional Review Board of the Charité Universitätsmedizin Berlin (EA4/006/22). Informed consent and consent to publish was obtained from each volunteer in accordance with the Declaration of Helsinki. All personal information collected from volunteers were treated with utmost confidentiality. For experiments on human macrophages, PBMC were isolated by performing a density centrifugation in SepMate tubes (Stem cell) using lymphoprep (Stem cell) density gradient medium. PBMC were then plated in DMEM culture medium supplemented with standard antibiotics, 10% FCS and human 50ng/ml GM-CSF (peprotech) at a density of 1×10^6^ cells/ml. The medium was changed every 2-3 days until the cells reached a full confluence.

### Non-canonical Inflammasome Induction in Human Macrophages

After 8 days of culture the medium was replaced by 50ng/ml GM-CSF containing 100µmol NAD^+^ culture medium. For 2 following days NAD^+^ was added daily until stimulation. To induce non-canonical inflammasome activation in human macrophages, cells were primed with 1µg/ml Pam3CSK4 for 5-6 hours. Subsequently the medium was replaced, and cells were treated with 3µg/ml LPS O111:B4 and 0.25% (vol/vol) Fugene HD Plus (Promega) to cause transfection. Finally, plates were centrifuged at 805 x g for 2 minutes and subsequently cultured for 20 hours at 37°C, 5% CO_2_.

### Western blot

For Western blot analysis, proteins were extracted using RIPA buffer and the concentrations determined using Pierce^™^ BCA Protein Assay Kit. Subsequently, proteins were resolved in SDS-PAGE, transferred to 0.45 μm nitrocellulose membranes (BioRad), blocked with 5% non-fat dry milk in PBS with 0.1% Tween 20, and processed for immunodetection. The following primary antibodies were used according to manufacturer’s instructions: Pro-Caspase-1 (#ab179515, Abcam), Caspase-1 (**#**14-9832-82, eBioscience) IL-1β (AF-401-NA, RD Systems), NLRP3 (#768319, RD Systems), Caspase-11 (#mab8648, RD Systems), Gasdermin D (ab209845, Abcam), P-STAT-1 (#9167S, Cell Signaling), STAT-1 (#9172S, Cell Signaling), NF-κB-p65 (#49445S, Cell Signaling), NF-κB-p52 (#4882S, Cell Signaling), β-Actin (ab3280, Abcam). Antibody detection was performed with HRP-coupled goat secondary anti-mouse or anti-rabbit antibodies (Immunoresearch), followed by ECL reaction (Perkin Elmer) and exposure to Fuji X-ray films. Finally, films were scanned, and signals quantified using the web-based image processing software ImageJ (NIH).

### Analysis of LPS-Transfection Efficiency

For intracellular detection of LPS, primed BMDMs were stimulated with 20µg/ml CTB and FITC-conjugated LPS O111:B4 for 16 hours, washed twice with PBS, fixed in 4% PFA containing PBS and DAPI for 10 minutes and subsequently analyzed using a confocal microscope and flow cytometry. To determine transfection efficiency using confocal microscopy, FITC-stained pixels per image were quantified using the web-based image processing software ImageJ (NIH).

### Caspase-1 assay

To determine the expression of Caspase-1, primed BMDMs were stimulated with 20µg/ml CTB and 2µg/ml LPS O111:B4 for 4 and 16 hours respectively, washed twice with PBS, stained using a caspase-1 active staining kit (Abcam) including caspase-1 staining (fluorescent green) and DAPI staining (fluorescent blue) according to the manufacturer’s protocol and analyzed using a confocal microscope.

### Endotoxic shock model

8-12-week-old C57BL/6 mice were treated with 40mg NAD^+^ for 2 following days before intraperitoneal injection of 54 mg/kg LPS O111:B4 or LPS O55:B5. Where indicated mice were administered 6mg/kg poly(I:C) 6 hours prior to LPS administration. Consequently, survival and body temperature were monitored every 2-4 hours for up to 100 hours. To assess the amount of systemic IL-1β and IL-18 by Elisa (both Invitrogen), mice were euthanized by decapitation 10 hours and 15 hours after LPS injection serum was isolated from collected blood.

### Flow cytometric analysis

To analyze splenic lymphocytes for the intracellular expression of IL-10 mice were challenged with 54 mg/kg LPS O111:B4 for 10 hours and euthanized by cervical dislocation subsequently. Spleens were harvested in a sterile environment and single cell suspensions were obtained using a 70um Nylon cell strainer.

Then, 1×10^6^ splenocytes per animal per condition were cultured in RPMI 1640 (#10-040-CV, Corning) supplemented with 10% BenchMark Fetal Bovine Serum (#100-106, Gemini), 1% penicillin/streptomycin (#30-002-CI, Corning), 2 mM L-glutamine (#25-005-CI, Corning), 20 ng/mL phorbol 12-myristate 13-acetate (PMA, #P8139-1MG, Sigma-Aldrich), 1 μg/mL ionomycin (#I9657-1MG, Sigma-Aldrich), and 0.67 μL/mL BD GolgiStop (#554724, BD Biosciences) for 4 hours at 37°C and 5% CO2 in 1 mL-volumes in a 12-well plate. After 4 hours, the cells were collected from each 12-well plate well and prepared for flow cytometry by staining the surface epitopes in flow staining buffer consisting of 1× DPBS supplemented with 1.0% (w/v) bovine serum albumin (#A2153, Sigma-Aldrich) and 0.020% sodium azide (#S8032, Sigma-Aldrich) for 25 min at 4°C. Then, the cells were fixed and permeabilized with the eBioscience Foxp3 Fixation/Permeabilization concentrate and diluent cocktail (#00-5523-00, Invitrogen) for 30 min at 4°C. Finally, the intracellular cytokine target was stained in 1× permeabilization buffer diluted from 10× eBioscience Foxp3 Permeabilization Buffer (#00-5523-00, Invitrogen) with deionized water. Finally, the stained samples were analyzed on a FACS Canto II (BD Biosciences, San Jose, CA, United States) flow cytometer, and the resultant flow cytometry standard (FCS) files were analyzed with FlowJo version 10 (Flowjo LLC, Ashland, OR, United States).

### Bacterial infection model

Frozen stock suspensions of Escherichia coli (Migul) (ATCC, 700928) were obtained from ATCC and cultured in 5ml Luria-Bertani medium at 37°C. Bacterial concentration was determined by plating 100ul, 10-fold serial diluted bacterial samples and counting the colony-forming units (CFU) after overnight incubation at 37°C. One day prior to injection 1ml of culture was reinoculated into 5ml of medium and incubated for 3-5 hours using a 37°C shaker at 250rpm agitation. Bacterial cultures were then centrifuged for 10 minutes at 3000rpm and washed twice with PBS. Mice were previously treated with NAD^+^ for 2 serial days and subsequently infected with E. coli by injecting 1ml of 1×10^9^ CFU/ml bacterial suspension intraperitoneally. The survival was monitored. In another set of experiments mice were sacrificed 5h hours after infection and kidneys and liver were harvested. The collected tissues were homogenized in 1ml of sterile PBS and 10-fold serial dilutions plated overnight at 37°C on LB agar plates to determine bacterial load per gram.

## Data Availability Statement

All data generated and analyzed during this study are provided with the manuscript as source data files for each figure. Data generated from RNA sequencing of BMDMs have been made publicly available in Dryad (DOI: 10.5061/dryad.zw3r228fj).

## Acknowledgements

J.I. was supported by the Berlin Institutes of Health Junior Clinician Scientist Program. Y.N. was supported by the Chinese Scholarship Council (201606370196) and Central South University. H.R.C.B. was supported by the Swiss Society of Cardiac Surgery. A.V. was supported by awards from the National Institute of Mental Health (R01MH110438) and National Institute of

Neurological Disorders and Stroke (R01NS100808).

---

**Figure 1 Supplement 1.**
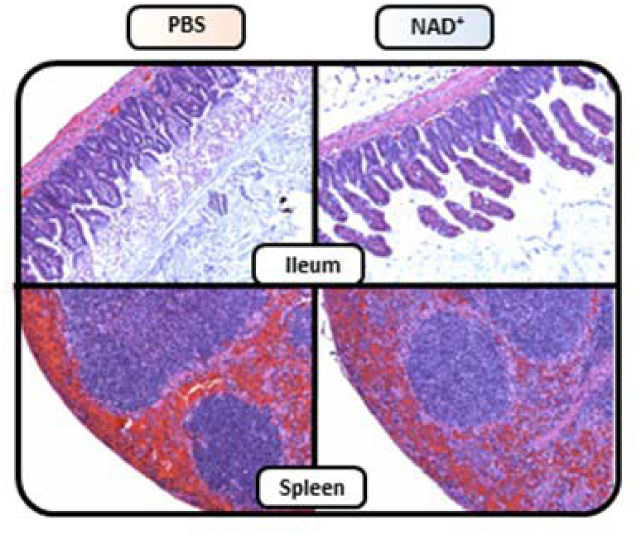
NAD^+^ preserves ileal villi structure and reduces splenic hemorrhage during LPS induced septic shock. C57BL/6 mice were treated with PBS or NAD^+^ for 2 days prior to administration of a lethal dose of LPS (O55:B5 /O111:B4) by intraperitoneal injection. Ileum and Spleen were removed after 15 hours and subsequently IHC for H&E staining was performed. All data depicted in this figure are provided as source data. Figure 1 Supplement 1 Source Data 1 – Raw Data for **Fig.1 Supplement 1**: Histology

**Figure 1 Supplement 2.**
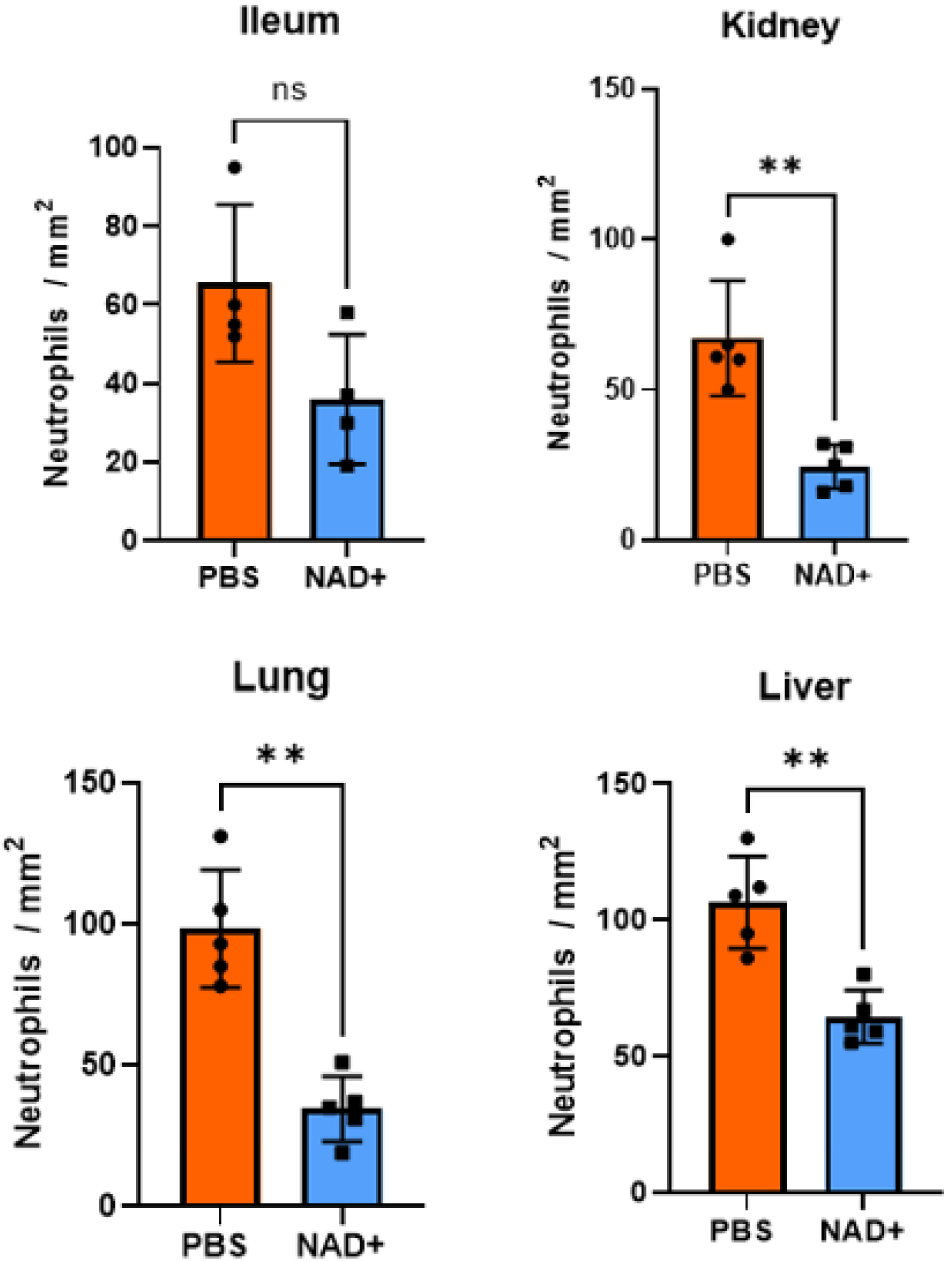
Neutrophils per mm^2^ infiltrating mice Ileum, Kidney Lung and Liver in the IHC stains. C57BL/6 mice were treated with PBS or NAD^+^ for 2 days prior to administration of a lethal dose of LPS (O55:B5 /O111:B4) by intraperitoneal injection. After a 15-hour interval, the ileums, lungs, kidneys, and livers were extracted and subjected to immunohistochemical staining with hematoxylin and eosin (H&E). Neutrophil quantification was performed subsequent to the immunohistochemical staining in various tissue samples. Notably, in the Kidney, Lung, and Liver tissues, the number of neutrophils was observed to be significantly higher in the PBS-treated mice group as compared to the NAD^+^ treated mice group. Column plots display mean with standard deviation (n=4-5). Statistical significance was determined by using Student’s T-test or One-Way-ANOVA. Asterisks indicate p-values * = p<0.05, **= p<0.01. All data depicted in this figure are provided as source data. Figure 1 Supplement 2 Source Data 1 – Raw Data for **Fig.1 Supplement 2**: Neutrophil Count

**Figure 2 Supplement 1.**
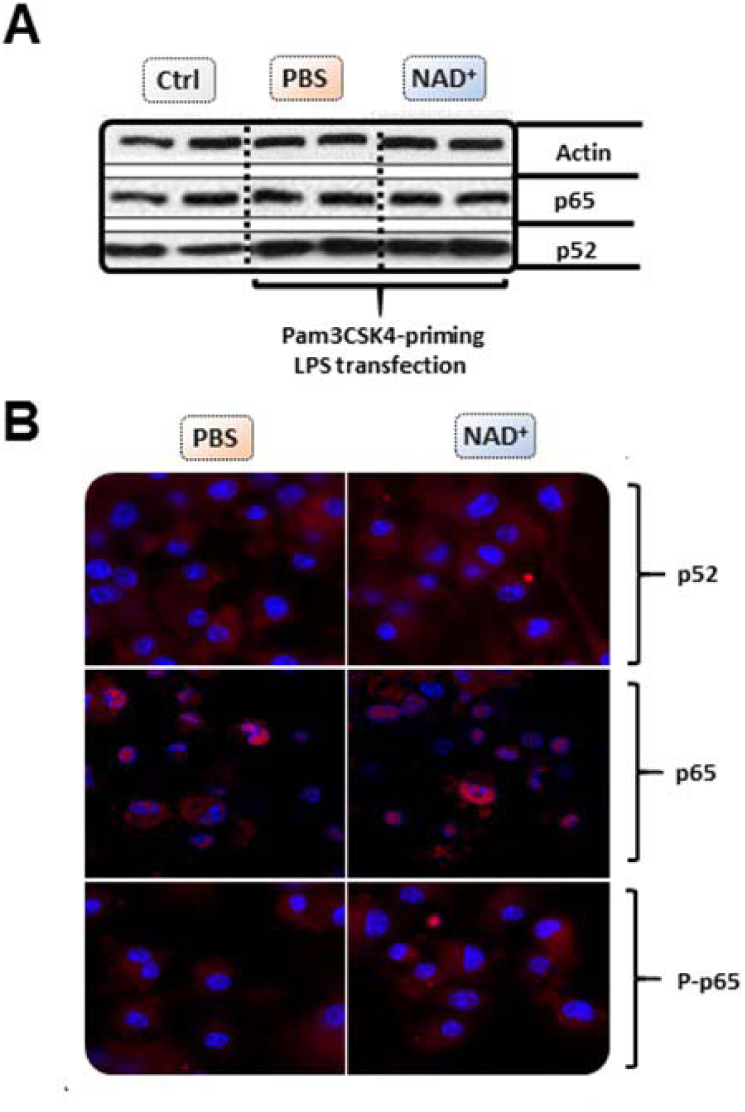
NAD^+^ does not alter BMDM derived NF-κB expression or phosphorylation. Differentiated BMDM were cultured in the presence of 100µmol NAD^+^ or PBS for 2 following days. BMDMs were then primed with 1µg/ml Pam3CSK4 and subsequently stimulated with 2µg/ml LPS O111:B4 and 20µg/ml CTB. Unstimulated BMDMs served as controls. (**A)** P52 and p65 expression was determined using Western blot. **(B)** stimulated BMDMs were stained with p52, p65 and phospho-p65 and expression levels assessed using confocal microscopy. All data depicted in this figure are provided as source data. Figure 2 Supplement 1 Source Data 1 – Raw Data for **Fig.2 Supplement 1 A**: Western Blot Figure 2 Supplement 1 Source Data 2 – Raw Data for **Fig.2 Supplement 1A**: Western Blots Bands Highlighted and Sample Labels Figure 2 Supplement 1 Source Data 3 – Raw Data for **Fig.2 Supplement 1 B**: Immunofluorescence

**Figure 2 Supplement 2.**
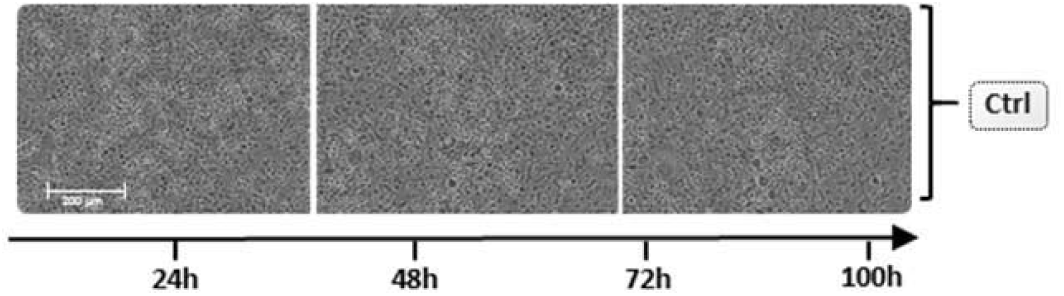
Unstimulated BMDM cell viability and apoptosis. Differentiated BMDMs were cultured in the presence of 100µmol NAD^+^ or PBS for 2 following days. BMDMs were then primed with 1µg/ml Pam3CSK4, subsequently stimulated with 2µg/ml LPS O111:B4 and 20µg/ml CTB and cell viability and apoptosis were monitored for 100 hours using the IncuCyte^®^ live microscopy system. All data depicted in this figure are provided as source data. Figure 2 Supplement 2 Source Data 1 – Raw Data for **Fig.2 Supplement 2**: Incucyte Live Microscopy

